# Identification and functional analysis of long non-coding RNAs in autism spectrum disorders

**DOI:** 10.1101/2020.03.15.986497

**Authors:** Zhan Tong, Yuan Zhou, Juan Wang

**Affiliations:** Department of Biomedical Informatics, School of Basic Medical Sciences, Peking University, Beijing 100191, China; Autism Research Center of Peking University Health Science Center, Beijing 100191, China

**Author notes:** To whom correspondence should be addressed: Juan Wang. E-mail for other co-authors: Zhan Tong and Yuan Zhou.

**Keywords:** long non-coding RNA, autism spectrum disorders, enrichment analysis, genome variants

## Abstract

**Background:** Genetic and environmental factors, alone or in combination, contribute to the pathogenesis of autism spectrum disorder (ASD). Although many protein-coding genes have now been identified as disease risk genes for ASD, a detailed illustration of long non-coding RNAs (lncRNAs) associated with ASD remains elusive. In this study, our aim was to identify ASD-related lncRNAs and explore their functions and associated biological pathways in autism.

**Methods:** ASD-related lncRNAs were identified based on genomic variant data of individuals with ASD from a twin study, and further validated using an independent copy number variant (CNV) dataset. The functions and associated biological pathways of ASD-related lncRNAs were explored by enrichment analysis of three different types of functional neighbor genes (i.e. genomic neighbors, competing endogenous RNA (ceRNA) neighbors and gene co-expression neighbors in the cortex). The differential functions of ASD-related lncRNAs in distinct brain regions were demonstrated by using gene co-expression network analysis based on tissue-specific gene expression profiles. Moreover, a functional network analysis were conducted for highly reliable functional neighbor genes of ASD-related lncRNAs. Finally, several potential drugs were predicted based on the enrichment of drug-induced pathway sets in ASD-altered biological pathway list.

**Results:** In total, 532 ASD-related lncRNAs were identified, and 86.7% of these ASD-related lncRNAs were further validated by a copy number variant (CNV) dataset. Most of functional neighbor genes of ASD-related lncRNAs were enriched in several functions and biological pathways, including nervous system development, inflammatory response and transcriptional regulation. As a set, ASD-related lncRNAs were mainly associated with nervous system development and dopaminergic synapse in the cortex, but associated with transcriptional regulation in the cerebellum. Moreover, all highly reliable functional neighbor genes were connected in a single functional network. Finally, several potential drugs were predicted and partly supported by the previous reports.

**Conclusions:** We concluded that ASD-related lncRNAs participate in the pathogenesis of ASD through various known biological pathways, which may be differential in distinct brain regions. And detailed investigation of ASD-related lncRNAs also provided clues for developing potential ASD diagnosis biomarker and therapy.

## BACKGROUND

Autism spectrum disorders (ASDs) are a group of heterogeneous neurodevelopmental disorders characterized by deficits in reciprocal social interaction and communication, and restricted interests and repetitive stereotypical behavior, with male-to-female prevalence nearly 3:1 [1, 2]. Approximately 10% of individuals with ASD have an identifiable genetic cause according to increasing clinical genetics services [3]. But because of highly genetic and phenotypic heterogeneity, the exact mechanism of ASD pathophysiology remains elusive [4-6]. Long non-coding RNAs (lncRNAs), which are defined as non-coding transcripts with more than 200 nucleotides in length, perform diverse regulation functions through a variety of mechanisms, including cell cycle regulation, RNA processing and editing, molecular scaffold, chromatin remodeling, genome imprinting, miRNA sponges and transcriptional regulation [7, 8]. It has been shown that the mutation and dysregulation of lncRNAs are involved in a wide variety of diseases, including cancer, neurological disorders and cardiovascular diseases [9-12]. And lncRNAs are also emerging as an important component in normal brain development [13]. Therefore, detail illustration of lncRNA and its function mechanism involved in ASD will greatly expand our understanding of the pathogenesis of ASD.

Twins and family studies have illustrated the predominant role of genetic factors in the pathogenesis of ASD [5, 6, 14]. Although many studies addressing the genetic architecture of ASD have mainly focused on illustrating the roles of protein-coding genes, increasing number of researchers are exploring the association between lncRNAs and the pathogenesis of ASD [6, 10, 15]. But most evidence, which characterized lncRNA dysregulation as an integral component of the transcriptomic signature of ASD, was derived from gene expression profiles of individuals with ASD [16-18]. Only several lncRNAs associated with ASD have been identified by genome-derived evidence, and further explored in action mechanisms by loss-of-function and gain-of-function experiments, such as *SHANK2-AS, MSNP1-AS* and *BDNF-AS* etc [19, 20]. In consideration of spatiotemporal-specific expression patterns, lncRNAs execute different functions and have unique gene expression patterns in distinct cellular conditions, therefore differentially expressed lncRNAs in a specific tissue couldn’t reflect the global effects of dysregulated lncRNAs [21]. Identification and functional analysis of genome-derived lncRNAs may complement these limitations.

Gene co-expression analysis and genomic neighbor region analysis are recognized as two traditional annotation ways of the biological functions and associated conditions of uncharacterized lncRNAs, which have a wide range of applications [22, 23]. Competing endogenous RNA (ceRNA) crosstalk, which refers to a hypothesis that all RNA transcripts can communicate with and regulate each other through competing to bind shared miRNAs, has been increasingly recognized as an important way to study gene functions and associated biological mechanisms [24]. In this study, we first identified ASD-related lncRNAs based on the genome variants detected in individuals with ASD from the previous twin study, which defined the discordant variations in monozygotic twin (DVMT) occurred in at least two twin pairs as putative ASD risk sites [25]. These ASD-related lncRNAs were found to have less gene essentiality scores and greater gene expression specificities, and tended to have greater median expression in the brain tissues when compared to all profiled tissues. Then we explored the functions and associated mechanisms of ASD-related lncRNAs using enrichment analysis of three distinct types of functional neighbor genes, including genomic neighbors, ceRNA neighbors and gene co-expression neighbors (derived from gene expression profiles in the cortex). And we also explored the distinct functions of ASD-related lncRNAs in different brain regions. Furthermore, a functional interaction network was constructed for the highly reliable functional neighbor genes of ASD-related lncRNAs. Finally, several drugs which have potential to intervene the aberrant state of ASD in the biological pathway level, were predicted.

## METHODS

### Identifying long non-coding RNAs (lncRNAs) in autism spectrum disorder

We obtained the genomic variant data of individuals with autism spectrum disorder (ASD) from Huang et al.’s study [25], which identified genomic variants in monozygotic twins discordant for ASD using genome-wide sequencing. Then the genomic coordinate data of reference genes from UCSC genome browser (https://genome.ucsc.edu/) [26] (i.e. known gene track) and Ensembl database (http://www.ensembl.org/) [27] were used to intersect the genomic variant data to identifying long non-coding RNAs (lncRNAs) with overlapping mutations, which were so-called ASD-related lncRNAs in the following.

To explore the reliability of ASD-related lncRNAs obtained from the twins’ study, we downloaded the copy number variation (CNV) dataset from the Simons Foundation Autism Research Initiative (SFARI) database (https://gene.sfari.org/autdb/GS_Home.do, SFARI-Gene_cnvs_06-20-2019release) [28]. All genomic coordinates of the CNV dataset were transferred to the latest genome assembly (hg38) using UCSC liftOver tool, and the regions that failed to converted were discarded. Then we established the CNV-related lncRNAs from this dataset using the pipeline described above.

### Gene importance and expression specificity analysis of ASD-related lncRNAs

We downloaded gene importance score data from Gene Importance Calculator (GIC) database (http://www.cuilab.cn/gic), which defining gene essentiality scores of protein-coding genes and lncRNAs based on sequence features [29]. Then the Wilcoxon rank sum test was performed to investigate GIC score characteristic of ASD-related lncRNAs.

The tissue-specific RNA-seq dataset was downloaded from the Genotype-Tissue Expression (GTEx) project (http://gtexportal.org, 2016-01-15_v7_RNASeQCv1.1.8 release) [30], in which gene-level median TPM values are reported for 53 different tissue types. Brain tissue expression index was calculated by the median of the expression values in brain tissues divided by the median expression for all tissue types. On the basis of the RNA-seq dataset, the tissue specificity index τ was calculated for each gene based on the program proposed by Yanai et al [31]. The Wilcoxon rank sum test was then performed to explore the brain tissue expression index and tissue specificity index characteristics of ASD-related lncRNAs.

### Weighted gene co-expression network analysis (WGCNA)

We constructed specific gene co-expression networks for different brain regions using the WGCNA package in R (v1.66) [32]. To filter many genes with subtle expression or limited expression variation across different samples, only genes with top 75% median absolute deviation (MAD) across samples were retained. All genes and samples were then checked for excessive missing values, and the obvious outliers were excluded by sample hierarchical clustering. The function *pickSoftThreshold* provided by the WGCNA package was used to choose the soft thresholding power to approximate network scale-free topology. Afterwards, an unsigned weighted network was created using the selected soft thresholding power and the Pearson correlation. Based on the gene expression profiles and associated sample annotation file from GTEx database (http://gtexportal.org, 2016-01-15_v7_RNASeQCv1.1.8 release), totally seven brain region specific gene co-expression networks (i.e. cortex, frontal cortex (BA9), anterior cingulate cortex (BA24), cerebellum, hippocampus, hypothalamus and amygdala) were constructed, respectively.

### Establishment of functional neighbor genes of ASD-related lncRNAs

According to the previous studies, there occur three complementary ways to predict the unknown function of lncRNA, including enrichment analysis of lncRNA’s genomic neighbors, competing endogenous RNAs (ceRNAs) neighbors and gene co-expression neighbors. In this study, we chose the upstream and downstream 50kb flanking region of lncRNA gene as its neighbor region [23]. Then BEDTools [33] was used to find the overlaps between the genomic coordinates of reference genes from Ensembl database and the lncRNAs’ neighbor regions, and finally identified the genomic neighbors of ASD-related lncRNAs. Competing endogenous RNAs (ceRNAs), which share at least two regulating miRNAs with ASD-related lncRNAs, were retrieved from starBase v2.0 database (http://starbase.sysu.edu.cn/) with the threshold of p-value and false discovery rate (FDR) corrected p-value ≤0.01 [34]. These connected protein-coding genes were considered as the ceRNA neighbors of ASD-related lncRNAs. We speculated that the human cortex was more relevant to the pathophysiology of ASD compared to the other brain regions, based on much more ASD researches using the cortex samples [6, 17, 35]. So we obtained gene co-expression neighbors of ASD-related lncRNAs from the weighted gene co-expression network based on gene expression profiles of the cortex samples. For each functional neighbor gene set, only protein-coding genes were retained for the following analyses.

### Functional analysis

For each functional neighbor gene set, we conducted gene ontology (GO) and KEGG pathway enrichment analysis using DAVID web server (https://david.ncifcrf.gov/) [36, 37]. For functional neighbor genes which occur in at least two among three functional neighbor categories (genomic neighbors, ceRNA neighbors, gene co-expression neighbors in the cortex), we explored the functional network of these genes using GeneMANIA (http://genemania.org/) [38]. Notably, a more stringent threshold for filtering gene co-expression neighbors was used to obtain a reasonable number of highly reliable functional neighbor genes in the analysis of functional network.

### Potential drug prediction

We downloaded the drug-induced KEGG pathway dataset from Drug-Path database (http://www.cuilab.cn/drugpath) [39], which predicted drug-induced pathways based on the transcriptome downloaded from the CMap database. Then we grouped the KEGG pathways by the same drugs to generate the specific drug-induced KEGG pathway sets. The fisher’s exact test was used to find the overrepresentation of each drug-induced KEGG pathway set (i.e. potential drug) in the altered KEGG pathway list obtained from functional analysis of ASD-related lncRNAs. To obtain the altered KEGG pathway list with appropriate number, we used FDR ≤ 0.5 as the filtering threshold. All human KEGG pathways included in the KEGG database (https://www.genome.jp/kegg/) [40] were taken as the background.

## RESULTS

### Identification and characterization of long non-coding RNAs in autism spectrum disorders

Firstly we downloaded the genomic variant dataset of individuals with autism spectrum disorders (ASDs) from the previous study [25], which included single nucleotide variants (SNVs), small insertions or deletions (Indels) and copy number variants (CNVs). Then the genomic locations of reference genes were scanned to identify ASD-related long non-coding RNAs (lncRNAs) which had overlapped mutations associated with ASD. Totally, 532 ASD-related lncRNAs were identified and further classified into several categories based on gene type annotations from Ensembl database (see Additional file 1: Table S1) [27]. We found that more than a half of ASD-related lncRNAs came from genome intergenic regions, and lncRNAs which were generated from the antisense strand of protein-coding genes, also occupied a large proportion (Figure 1A). Since the original genome variant dataset was derived from small samples and single cohort, we also validated these ASD-related lncRNAs with the CNV dataset from SFARI database [28]. 86.7% of ASD-related lncRNAs were shown to overlap with at least one ASD-associated CNV. Through the characteristics analyses, ASD-related lncRNAs were shown to have lower gene importance, higher tissue expression specificity compared to the background genes (Wilcoxon rank sum test, P-value = 1.61E-18 and 2.79E-13, respectively) (Figure 1B-C). Moreover, ASD-related lncRNAs seem to have bigger brain tissue expression index (i.e. the ratio of median gene expression in brain tissues vs all profiled tissues) compared to the background, but no significant difference was detected (Figure 1D).

**Figure 1.**
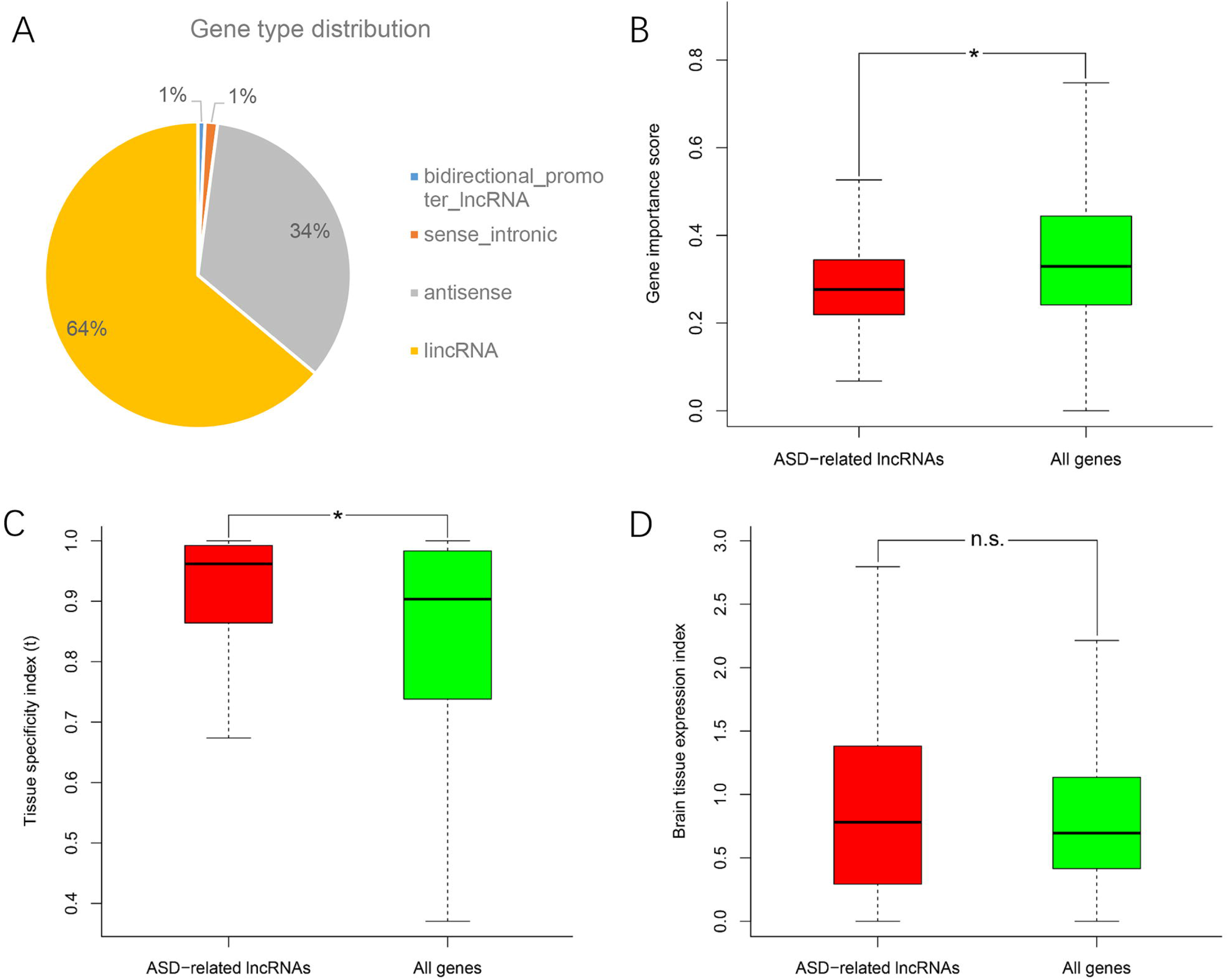
Gene type distribution and characteristics analysis of ASD-related lncRNAs. (A) Pie chart displays the distribution of ASD-related lncRNAs in different gene types. Comparison of ASD-related lncRNAs with the background genes in (B) gene importance score, (C) tissue specificity index τ and (D) brain tissue expression index. *P-value < 10e-5 from Wilcoxon test. N.s. represents no statistical significance.

### Functional enrichment analysis of ASD-related lncRNAs

To characterize the functions and associated biological pathways of ASD-related lncRNAs, it is an important step to find their functional-relevant protein-coding genes. LncRNAs are recognized as potential cis-regulators of their genomic neighbor genes [41], so it is a reasonable way to predict the functions and implicated biological pathways of ASD-related lncRNAs based on their genomic neighbors. Gene ontology (GO) enrichment analysis of genomic neighbors has shown that most of these genes were enriched in a series of immune pathways, including cellular response to interferon-gamma, cellular response to interleukin-1 and positive regulation of inflammatory response (Figure 2A) [42]. KEGG pathway analyses illustrated the enrichment in tyrosine metabolism pathway, which was consistent with the fact that MET receptor tyrosine kinase (*RTK*) is an autism risk factor (Figure 2B) [43]. Together, these results indicate that ASD-related lncRNAs may be implicated in the pathogenesis of autism partly through immune response processes.

**Figure 2.**
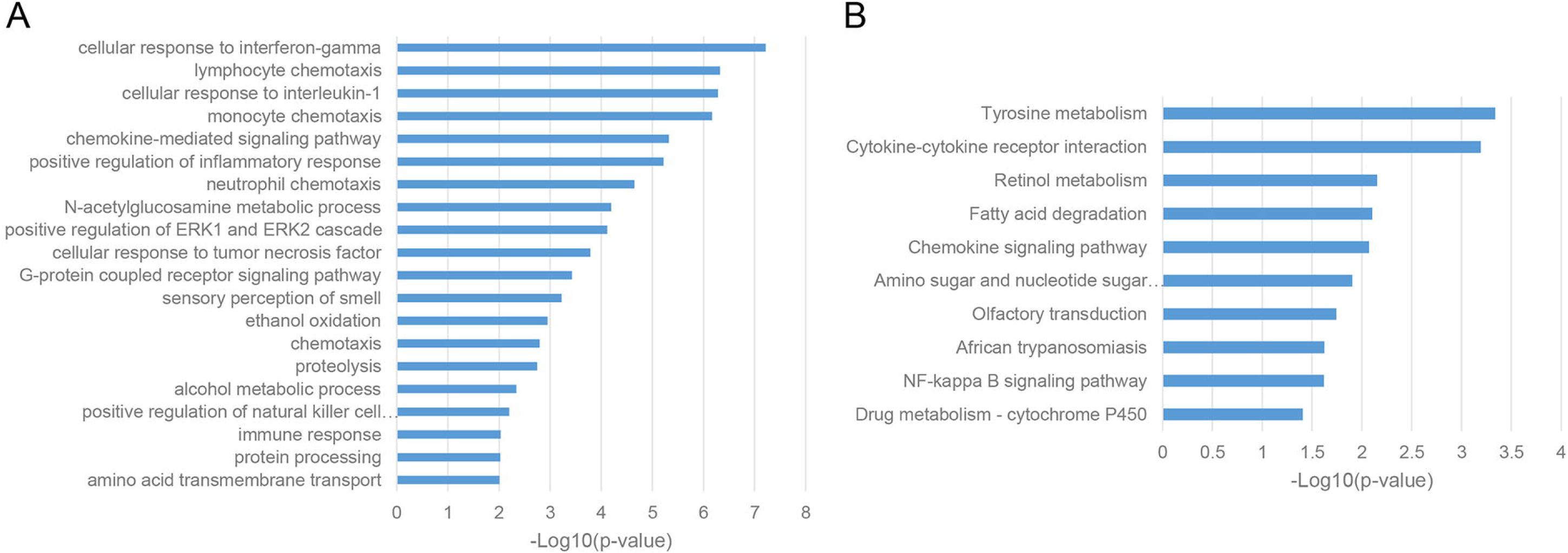
Enriched functional terms of genomic neighbors of ASD-related lncRNAs. (A) Top 20 gene ontology biological process (GO BP) terms with p-value < 0.05. (B) KEGG pathway terms with p-value < 0.05.

Given the fact that lncRNAs can regulate protein-coding mRNAs through the mechanism of miRNA sponges [24], we conducted the enrichment analysis of competing endogenous RNA (ceRNA) neighbors of ASD-related lncRNAs. We found that most of these genes were enriched in transcriptional regulation, such as positive regulation of transcription from RNA polymerase II promoter and positive regulation of transcription, DNA-templated (Figure 3A) [44]. Furthermore, KEGG pathway analyses uncovered the implication of insulin signaling pathway, Glioma and Wnt signaling pathway, which was consistent with the previous reports (Figure 3B) [5, 6].

**Figure 3.**
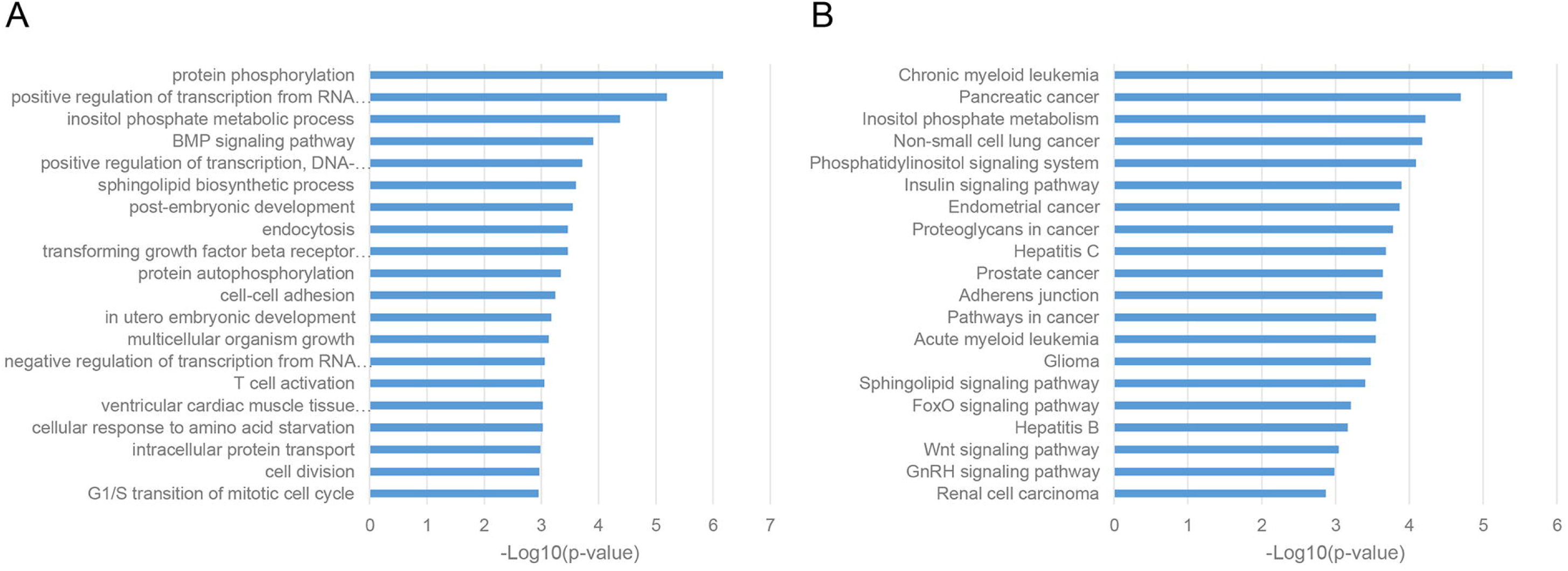
Enriched functional terms of competing endogenous RNA (ceRNA) neighbors of ASD-related lncRNAs. (A) Top 20 GO BP terms with p-value < 0.05. (B) Top 20 KEGG pathway terms with p-value < 0.05.

As a classic way for predicting the function of uncharacterized genes, we also performed the enrichment analysis of gene co-expression neighbors of ASD-related lncRNAs in human cortex, which was widely accepted as the most relevant tissue in the pathogenesis of ASD. As shown in Figure 4A-B, gene co-expression neighbors have shown the most relevance with the neuropathological mechanisms of ASD among these three categories of functional neighbor genes. Most of gene co-expression neighbors were enriched in nervous system associated biological processes, such as chemical synaptic transmission, neurotransmitter secretion and nervous system development from GO biological process terms, synaptic vesicle cycle and dopaminergic synapse from KEGG pathways [21, 25, 44]. Notably, MAPK signaling pathway was identified by both enrichment analysis categories, which were inferred to involve in the pathogenesis of ASD through the regulation of cell-proliferation pathways in brain development [5].

**Figure 4.**
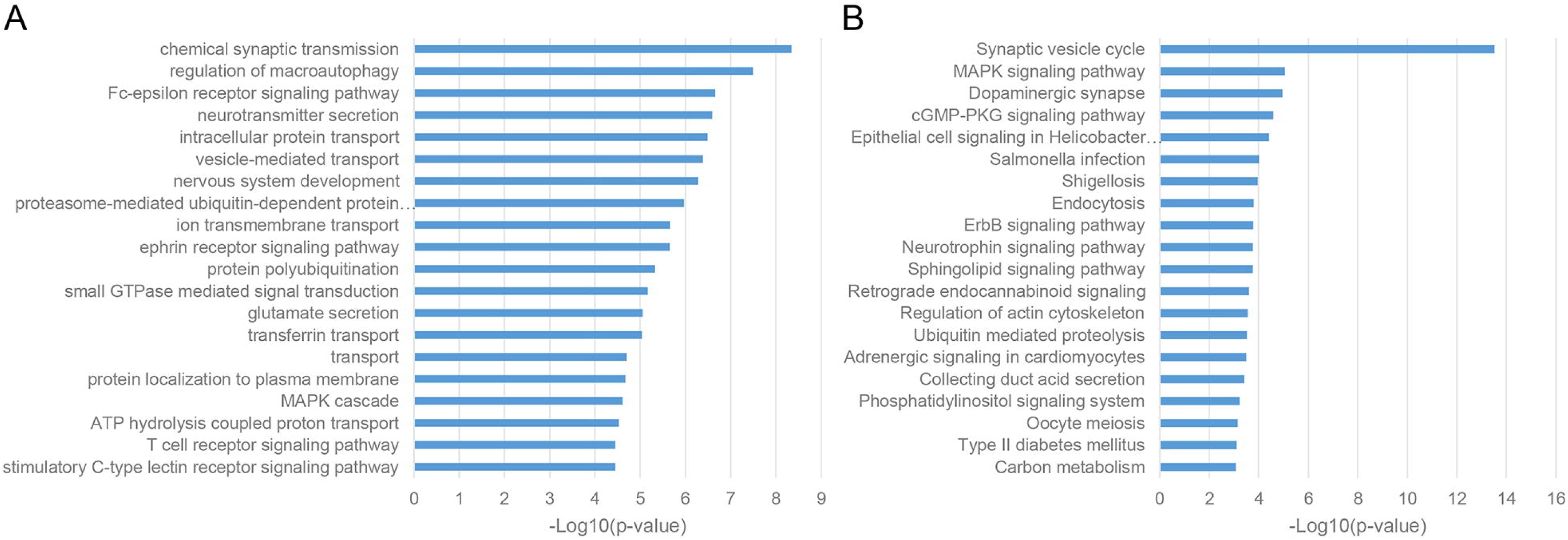
Enriched functional terms of gene co-expression neighbors of ASD-related lncRNAs. (A) Top 20 GO BP terms with p-value < 0.05. (B) Top 20 KEGG pathway terms with p-value < 0.05.

All these results provided the evidences of involvement of ASD-related lncRNAs in the pathogenesis of autism. It also implied that ASD-related lncRNAs may participate in various ASD implicated biological pathways through multiple regulation mechanisms when considering the complementary enrichment analysis results of different types of functional neighbor genes. These results also prompt some additional biological pathways potentially associated with ASD. For example, proteolysis and/or protein ubiquitination were uncovered in the enrichment analysis of both genomic and co-expression neighbors, but the detailed illustration of its involvement in autism remains unclear.

### Differential enriched functional terms of ASD-related lncRNAs in different brain regions

During the gene co-expression analysis in human cortex, we noticed that it was interesting to illustrate whether ASD-related lncRNAs executed similar functions in different brain regions. Several brain regions have been reported to be associated with the pathogenesis of ASD, such as cortex, frontal cortex (BA9), anterior cingulate cortex (BA24), cerebellum, hippocampus, hypothalamus and amygdala [6, 43, 45, 46]. So we constructed specific weighted gene co-expression networks for these distinct brain regions, and then performed enrichment analysis using the pipeline described before, respectively. Actually, the results of enrichment analysis have shown the differential functions of ASD-related lncRNAs as a class in distinct brain regions (Figure 5A-B). Most of gene co-expression neighbors of ASD-related lncRNAs in the cerebellum were enriched in transcriptional regulation and DNA repair, while neighbors in the cortex were enriched in a plenty of nervous system related functions, in consistence with the important roles of cortex in the pathogenesis of ASD. Interestingly, neighbors in BA9 and BA24 have shown significant differences in enriched function terms. Most of co-expression neighbors in BA9 were enriched in protein ubiquitination and ubiquitin mediated proteolysis, while neighbors in BA24 have also shown enrichment in several neuron related functions, including myelination, axonogenesis and etc. Previous studies have illustrated the excess of 67% neurons in the prefrontal, and a quantitative gradient of brain overgrowth from anterior/frontal to posterior in the majority of individuals with ASD [21]. One plausible mechanism would be the degree of dysregulation of ubiquitin-proteasome dependent degradation resulting in the gradient of neural cell growth [47]. Furthermore, most of gene co-expression neighbors in the hippocampus and hypothalamus were also enriched in several nervous system functions, while neighbors in amygdala were enriched in cell adhesion and inflammatory response [5, 6, 21, 42].

**Figure 5.**
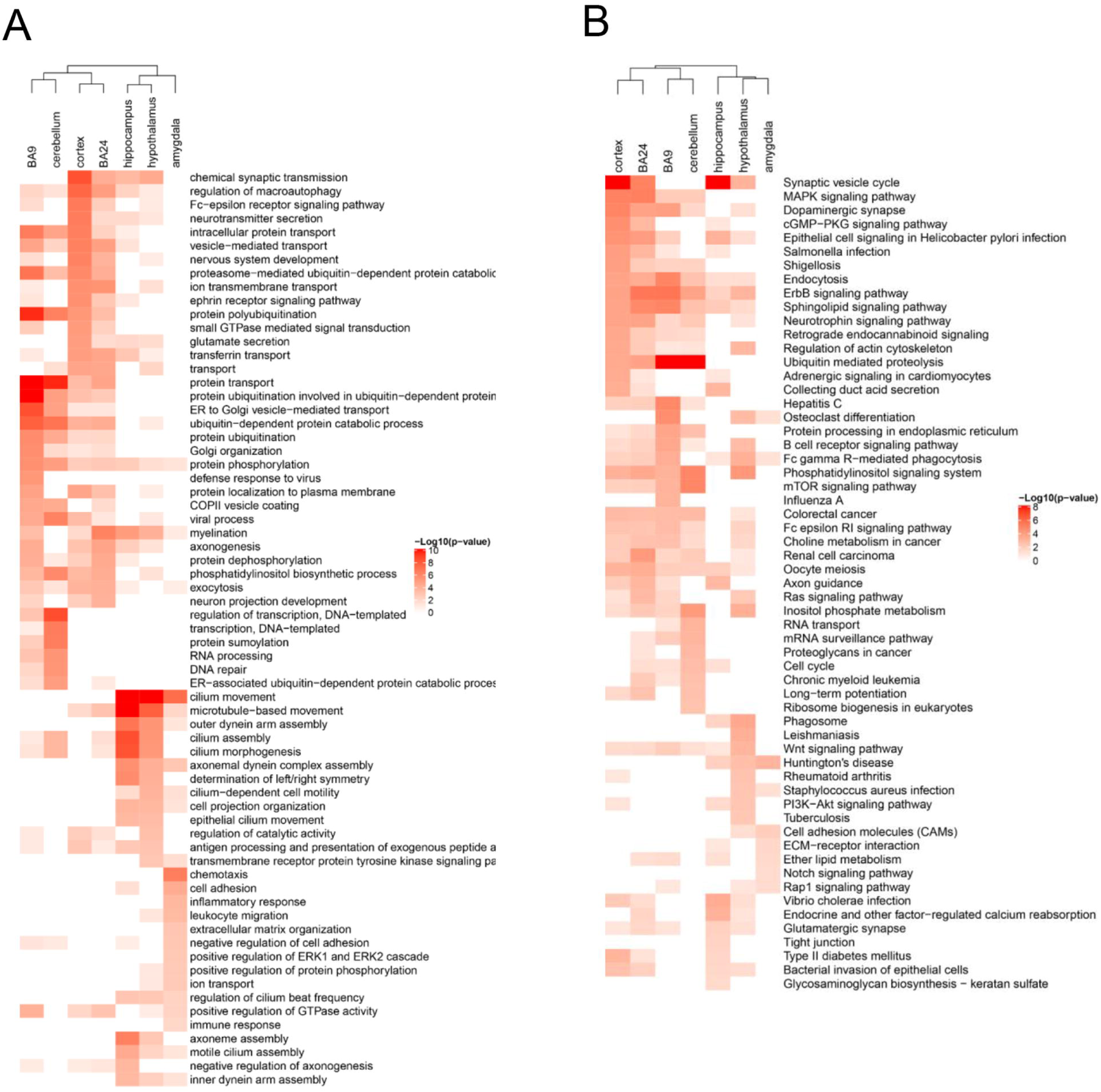
Differential enriched functional terms of ASD-related lncRNAs in distinct brain regions. Heatmaps display enriched (A) GO BP terms and (B) KEGG pathways of gene co-expression neighbors of ASD-related lncRNAs in different brain regions. Only top 15 terms with p-value < 0.05 in each brain region were depicted for both functional categories.

### Network analysis of highly reliable functional neighbor genes of ASD-related lncRNAs

To obtain the highly reliable functional neighbor genes of ASD-related lncRNAs, we intersected the three types of functional neighbor genes (i.e. genomic neighbors, ceRNA neighbors and gene co-expression neighbors filtered with the more stringent threshold in the cortex), and only retained the functional neighbor genes that occurred in at least two of these three categories. Using GeneMANIA webserver, we found that all these highly reliable functional neighbor genes were connected to each other in a single functional network, which included 59.6% co-expression, 21.9% physical interaction, 9.0% co-localization, 5.0% pathway and 4.4% genetic interaction links (Figure 6B). And in this functional network, several genes were known to be associated with ASD, including *SYT1, DDX11, AGAP2, SLC4A10* and *SYNJ1* [28]. Moreover, these genes were related to several biological pathways, such as neurotransmitter secretion and transport, regulation of neurotransmitter levels, MAP kinase activity and etc.

**Figure 6.**
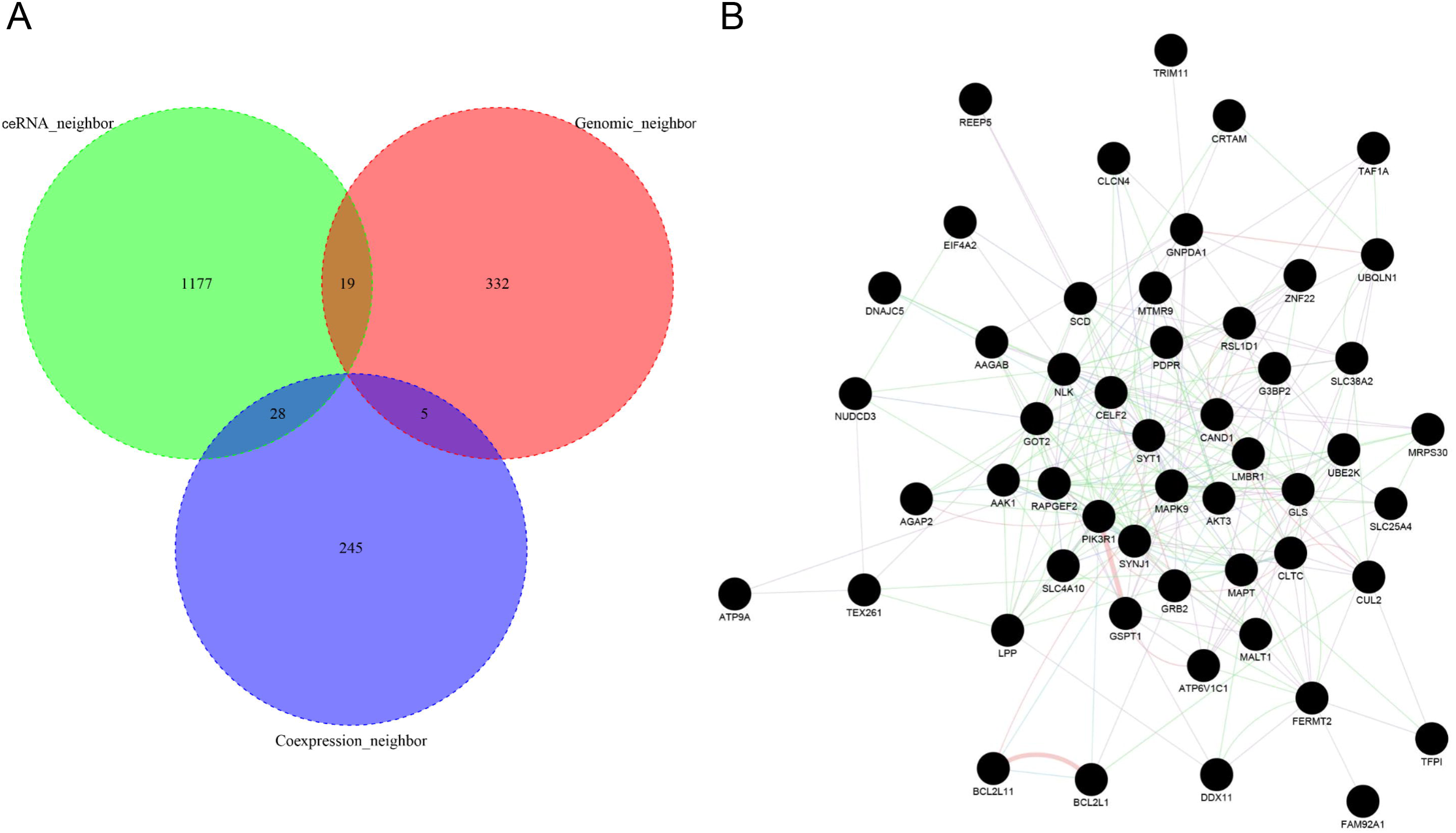
Functional network analysis of high-reliable functional neighbor genes of ASD-related lncRNAs. (A) Venn plot displays the intersection of three types of functional neighbor genes, i.e. genomic neighbors, competing endogenous RNA (ceRNA) neighbors and gene co-expression neighbors. (B) Network depicts the functional interaction relationships between each high-reliable functional neighbor gene pair.

### Potential drugs for intervening aberrant biological pathways in ASD

Take in consideration that heterogeneous genetic mutations of ASD were illustrated to converge in the common biological pathways [4, 6], we wondered whether there occurred some drugs which had potential to intervene the aberrant state of ASD in the biological pathway level. So we used fisher’s exact test to measure the overrepresented drug-induced KEGG pathway set (i.e. potential drug) in altered KEGG pathway list obtained from the functional analysis of ASD-related lncRNAs. Several enriched drugs predicted are potentially associated with ASD (Table 1). For example, amoxapine clinically used for depression treatment, has been demonstrated to have treatment benefits for interfering behaviors in individuals with ASD (false discovery rate (FDR) adjusted p-value = 0.0236) [48]. Piperine was experimentally supported to have abilities to reduce oxidative stress, elevate brain glutathione and improve behavior in VPA (valproic acid) -induced ASD mice model [49]. Diflunisal was shown to reduce the progression of neurological impairment and maintain quality of life in a randomized clinical trial for familial amyloid polyneuropathy [50]. Eticlopride, which was firstly developed for the treatment of schizophrenia, now is used in the study of the roles of D2-like receptor in schizophrenia and other brain diseases [51]. To explore the drug action targets of these potential drugs, we conducted the enrichment analysis with the DrugPattern tool by using these ten enriched drugs as input [52]. We found that the associated drug targets mainly included dopamine receptor, sodium-dependent serotonin transporter, adrenergic receptor, 5-hydroxytryptamine receptor and etc.

**Table 1.** Top 10 enriched drugs for potentially intervening aberrant KEGG pathways in ASD. P-values were calculated with fisher’s exact test. FDR represents the p-values adjusted by false discovery rate correction. The description information was retrieved from the KEGG drug [40], DrugBank [53], and PubChem [54] database.

| Drug name | P-value | FDR | Description |
| --- | --- | --- | --- |
| diflunisal | 9.62e-05 | 0.0236 | Analgesic, Anti-inflammatory, COX inhibitor |
| amoxapine | 8.91e-05 | 0.0236 | Antidepressant, Serotonin-noradrenaline reuptake inhibitor |
| budesonide | 3.29e-04 | 0.0605 | Antiasthmatic, Anti-inflammatory, Glucocorticoid receptor agonist |
| iopromide | 4.97e-04 | 0.0649 | Diagnostic aid (radiopaque medium) |
| PNU-0293363 | 7.04e-04 | 0.0649 | Not found |
| cyclizine | 6.86e-04 | 0.0649 | Anti-emetic, H1 receptor antagonist |
| ethoxyquin | 6.66e-04 | 0.0649 | UDP-glucuronosyltransferase activator, neuroprotective agent, Hsp90 inhibitor, genotoxin, food antioxidant. |
| piperine | 1.14e-03 | 0.0843 | NF-kappaB inhibitor |
| eticlopride | 1.05e-03 | 0.0843 | Dopamine Antagonists, Neurotransmitter Agents |
| nomegestrol | 1.32e-03 | 0.0882 | Ovulation inducing agent, Progesterone receptor agonist |

## DISCUSSION

LncRNAs, which constitute a large proportion of transcriptome, are increasingly recognized as the integral component of many fundamental biological processes and various disease pathogenesis. However, to our knowledge, limited efforts have been made to systematically characterize the functions and associated biological pathways of genome-level ASD-related lncRNAs. In this study, we first identified ASD-related lncRNAs using genome variant data in individuals with ASD downloaded from the previous study. Then the enrichment analysis of three types of functional neighbor genes provides abundant evidences of involvement of ASD-related lncRNAs in the pathogenesis of autism through various biological pathways, including nervous system development, chemical synaptic transmission, transcriptional regulation, immune pathway and etc. Functional network analysis of highly reliable functional neighbor genes of ASD-related lncRNAs (see more in Methods), which depicted a connected component, further implied that these ASD-related lncRNAs participated in the common biological pathways through possible gene expression regulation mechanism. Moreover, based on these altered KEGG pathways obtained from functional analysis of ASD-related lncRNAs, we predicted drugs with potential to intervene the disease status in the biological pathway level. Several interesting drugs were uncovered, such as amoxapine, diflunisal, budesonide, piperine and eticlopride. However, the mechanism of these drugs for the treatment of ASD still waited to be further experimentally explored.

In the meanwhile, multiple brain regions have been shown to be involved in the pathogenesis of autism. Interestingly, gene co-expression analysis in each brain region also revealed the differential functions and associated biological pathways of ASD-related lncRNAs in distinct brain regions. As illustrated by the enrichment analysis, co-expression neighbors in the cortex tended to be mostly enriched in nervous system related functions, while co-expression neighbors in the cerebellum were enriched in transcription regulation and DNA repair. And gene co-expression neighbors in frontal cortex (BA9) have shown more enrichment in ubiquitin mediated proteolysis when compared to those in anterior cingulate cortex (BA24). And most of gene co-expression neighbors in amygdala were enriched in cell adhesion and inflammatory response. All these results provide insight for studying differential function of ASD-related lncRNAs in distinct brain regions.

Though these results greatly expand our understanding of roles of lncRNAs in the genetic architecture of ASD, there are also some limitations in this study. At first, the genome variant data, which was downloaded from Huang et al.’s study [25], only included three pairs of monozygotic twins discordant for ASD. Statistical associations of these genome variants with ASD have not been explored properly and the cohort studied was single, which may cause possible errors in the detection of ASD-related lncRNAs or restrict the translation of the results to other cohorts. Moreover, we still lack evidences from transcriptome to validate the results we obtained based on the genome alternations. We believe that the transcript-level evidence could bridge genomic variants to transcriptome consequences, and provide additional clues for finding chemical molecular which could reverse the adverse transition in autism.

## CONCLUSIONS

In summary, we identified ASD-related lncRNAs and comprehensively explored their functions and associated biological pathways. We found that ASD-related lncRNAs participated in the pathogenesis of ASD through various biological pathways, which may also be different in distinct brain regions. Through functional network analysis, the highly reliable functional neighbor genes were shown to be connected in a single component, which further implied that ASD-related lncRNAs participated in the common biological pathways, such as neurotransmitter secretion and transport, regulation of neurotransmitter levels, MAP kinase activity and etc. Finally, several drugs which had potential to intervene the disease status of ASD in the biological pathway level, were predicted. In a word, taking lncRNAs into the framework of genetic architecture of ASD could draw a more comprehensive landscape of genetic factors and their interplays, and provide new approaches for ASD diagnosis and therapy.

## DECLARATIONS

### Ethics approval and consent to participate

Not applicable.

### Consent for publication

Not applicable.

### Availability of data and materials

The datasets used and/or analysed during the current study are available from the corresponding author on reasonable request.

### Authors’ contributions

J.W. conceived and designed this study. Y.Z. provided valuable comments and suggestions. Z.T. performed the analysis and drafted the manuscript. J.W. revised the manuscript. All authors read and approved the final manuscript.

## Acknowledgements

We thank the kindly sharing of the genomic variant data from Huang et al.’s study, which makes our identification and further analysis of the ASD-related lncRNAs become possible.

